# Jellyfish blooms Through the Microbial Lens: Temporal Changes, Cross-Species and Jellyfish-Water Comparisons

**DOI:** 10.1101/2024.05.23.595640

**Authors:** Noga Barak, Vera Brekhman, Dikla Aharonovich, Tamar Lotan, Daniel Sher

**Affiliations:** Department of Marine Biology, The Leon H. Charney School of Marine Sciences, University of Haifa, Haifa, Israel

**Author notes:** The fourth Author and the Fifth Author are co-senior authors.

**Keywords:** *Rhopilema nomadica*, 16S analysis, Microbiome, Cnidaria, bloom, jellyfish

## Abstract

In this study, we explore the dynamics of bacterial communities associated with *Rhopilema nomadica* blooms, the predominant jellyfish in the Eastern Mediterranean Sea. We collected over 120 samples from more than 30 individuals across five major bloom events, capturing both lesser-studied winter blooms and the peaks and declines of summer blooms. Our analysis revealed significant microbial shifts-increases in *Endozoicomonas* and unclassified Rickettsiales were significantly more abundance during late summer blooms, while *Tenacibaculum* dominated in winter. Additionally, we examined microbial patterns within specific tissues—bell, gonads, tentacles, and gastrovascular system—to assess variations across these different niches. This revealed high relative abundance of specific taxa tailored to different tissue-*Bacteroides* was predominantly found in the bell, Simkaniaceae in the gonads, and *Endozoicomonas* in the tentacles. Further expanding our research, we compared the top taxa of *R. nomadica* with those of nine other jellyfish species from different locations. Interestingly, while no universal core microbiome was found, several taxa, including *Endozoicomonas*, Mycoplasma, and *Spiroplasma*, were common across different species, suggesting their potential ecological roles across jellyfish. Lastly, our study of potential bacterial transmission modes revealed that key bacteria associated with *R. nomadica* are exclusively found near bloom areas, and are absent from remote seawater, highlighting potential localized transmission dynamics between jellyfish and their immediate marine environment. Our study marks the first exploration of microbial dynamics within *R. nomadica,* while also broadening the understanding of other jellyfish microbial communities and setting the stage for future studies to delve deeper into their complex interactions.

**IMPORTANCE:** Jellyfish blooms, like those of *Rhopilema nomadica* in the Eastern Mediterranean, impact marine ecosystems and human industries. Understanding the complex relationships between jellyfish and their microbiomes is important, as these interactions may influence bloom formation and decline. Our study explores microbiome variations across different stages of *R. nomadica* blooms, identifies common bacteria among jellyfish from various locations, and examines potential transmission modes of the main jellyfish-associated bacteria. Microbial communities vary significantly between bloom stages and jellyfish tissues, becoming less diverse towards the end of the bloom. Although no universal core microbiome was discovered, taxa such as *Endozoicomonas*, *Mycoplasma*, and *Spiroplasma* are prevalent across various jellyfish, suggesting significant ecological roles. Finally, our findings indicate that key bacteria associated with *R. nomadica* predominantly reside near bloom areas and are absent from distant seawater, highlighting localized transmission mode. This study enhances our understanding of jellyfish-associated microbial communities and their role in bloom dynamics.

## INTRODUCTION

Jellyfish blooms are a widespread and dynamic global phenomenon with broad ecological and economic implications. The effects of such blooms include damage to tourism, clogging of water intake of power stations and desalination plants, the disruption of local fisheries and, potentially, the alteration of marine food webs (1–4). These usually ephemeral jellyfish blooms, characterized by significant, often seasonal increases in jellyfish populations, are caused by a complex interplay of environmental factors and human activities (5–8). Typically, these blooms are more common during mid-spring until mid-autumn (8, 9) decline rapidly, leading at times to “jelly-falls”, where large amounts of biomass are exported to the seafloor (10–12). Thus, despite their frequent occurrence and substantial ecological and societal impact, the reasons why blooms appear and decline rapidly remain elusive.

One potential factor in jellyfish blooms is the interplay between jellyfish and their associated bacteria, viewing the bloom-forming jellyfish as a holobiont—a symbiotic unit of the jellyfish and its bacteria, or microbiome. This symbiosis can be crucial for the emergence of blooms, as evidenced by recent studies showing that bacteria are involved in various stages of early jellyfish development, such as larva settlement and the initiation of the strobilation process (7–9). As the jellyfish mature, these bacteria are hypothesized to play essential roles in nutrient acquisition, digestion, and immune response support (10–12). However, the disappearance of jellyfish blooms could be triggered by massive mortality events, potentially due to physiological changes associated with natural aging processes (5) or the introduction of harmful pathogens. Such events likely cause significant shifts in the microbial community associated with the jellyfish. These bacteria may be specifically adapted to different parts of the jellyfish or various stages of its lifecycle (13), each serving specialized roles. Alternatively, some may be opportunistic, thriving in the jellyfish environment without a definitive symbiotic relationship. These complex interactions highlight the need to closely examine the jellyfish microbiome throughout different stages of blooms to discern the microbial compositions that might influence the onset and decline of these events.

In the Eastern Mediterranean Sea, *R. nomadica* stands out as the predominant jellyfish species, known for forming significant blooms. First documented in the Mediterranean in 1977, this species has seen a notable population increase during the 1980s. It was initially observed off the coast of Israel, then spread along the Eastern Mediterranean coastline, reaching as far west as Sardinia and Sicily (14–19). The blooms appear predominantly during the summer months, with secondary blooms occurring in winter (14, 19). Here, we asked whether the *R. nomadica* microbiome is different between the winter and summer blooms, and whether it changes during summer between early bloom (June) and later stages, towards bloom disappearance (July). To answer this question, we also needed to take into account other factors that could affect the microbiome structure such as jellyfish size, sex, different tissues and jellyfish health. With this dataset, we then asked whether different types of jellyfish, each with their unique characteristics and backgrounds, host common bacteria (defined as having identical 16S rRNA sequences). This exploration could uncover bacteria essential across various jellyfish. Finally, we asked whether the main bacteria in the jellyfish can be identified in the water surrounding the bloom, as well as in an annual time-series from a reference location, where jellyfish were not observed at the time of sampling (the SoMMoS cruise series (20). This could help answer the question “how did the jellyfish get its bacteria?”.

## MATERIALS AND METHODS

### Sample Collection and DNA Extraction

A total of 31 jellyfish were collected during different bloom events around and in Haifa Bay, Israel (Fig. 1A-C). Two winter blooms were sampled (February 2020 and 2021), and a single summer bloom was sampled twice during the peak of the bloom in mid-June and toward the end of the bloom during mid-July. Each jellyfish was carefully collected into an 80L container filled with ambient seawater (21). The jellyfish were dissected on shore immediately following each research cruise (typically at most 2 hours after collection). To mitigate the risk of cross-contamination, a separate sterile toolkit was utilized for each jellyfish, with rigorous cleaning procedures implemented between the handling of different tissues. These included cleaning the tools with 1% sodium hypochlorite, DNA AWAY™ (Thermo Fisher Scientific), 70% ethanol, and ultrapure water after each use. For each jellyfish, three replicates of four distinct tissues (bell, gonads, tentacles, and gastrovascular canals (GVC)) were collected into sterile 0.5 ml Eppendorf tubes and immediately frozen on dry ice. Alongside tissue collection, the diameter of each jellyfish was measured, any unique characteristics were recorded, and a portion of the gonads was collected for sex determination in the lab. Additionally, at each location, three replicates of five litters of seawater were filtered through a Sterivex filter cartridge (0.22 μm), 1ml of preservation/lysis solution was added (40mM EDTA, 50mM Tris pH 8.3, 0.75M Sucrose), and the filters were frozen on dry ice. Upon arrival at the lab, the samples were transferred to a ^-^80°C freezer for long-term storage until further analysis.

**FIG 1:**
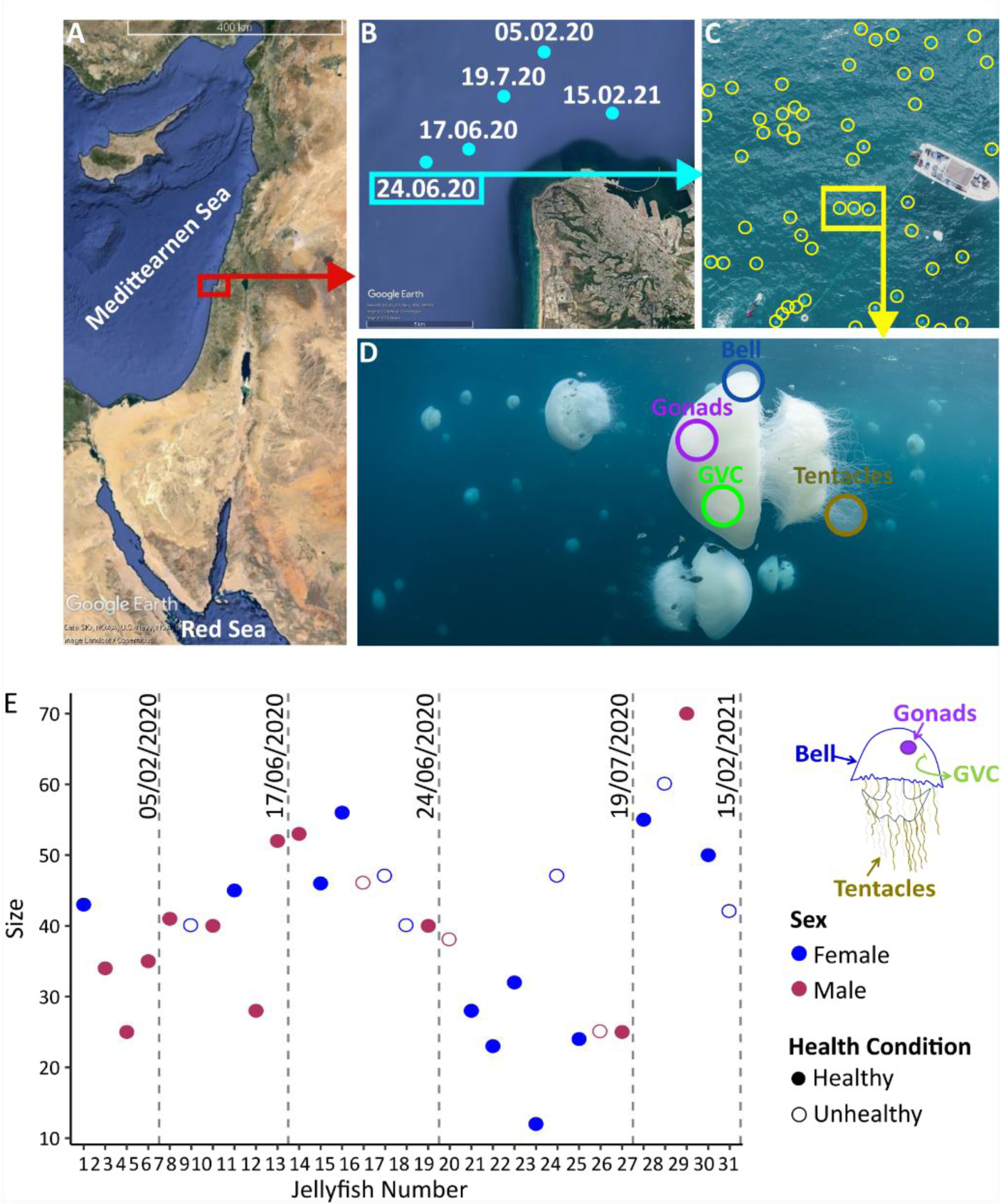
Jellyfish Sample Collection Overview. A, B) Maps of the sampling locations and the dates of sampling. C) A photograph taken from a drone deployed during the jellyfish sampling on 24.6.2020 (23), demonstrating the density of the jellyfish bloom. Individual jellyfish are marked by yellow circles. D) Underwater bloom perspective of a jellyfish bloom. Circles indicate the different tissues sampled from the jellyfish. Photo by Hagai Nativ. E) Collection data graph displays size, sex, health condition, and collection date for the 31 jellyfish. When jellyfish were identified as “unhealthy”, e.g. exhibited visible lesions on the bell, samples were collected from both visibly unhealthy bell tissue and healthy tissue from the same individual.

Samples from the bell, gonads, tentacles and GVC were thawed, homogenized using a bead beater (TissueLyser II, Qiagen), treated with 100 mg/ml lysozyme at 37°C (Merck) for one hour, and then incubated for an additional hour at 55°C after adding 20 mg/ml proteinase K (Promega). DNA was extracted using the ZymoBIOMICS DNA Miniprep Kit (Zymo Research), according to the manufacturer’s protocol. The Sterivex filter (Millipore) was processed as described previoulsy (22), with the filter being placed in the ZR BashingBead™ lysis tubes from the ZymoBIOMICS DNA Miniprep Kit. The subsequent DNA extraction was carried out as described for the tissue samples. For standardization purposes, a commercially available mock microbial community standard (ZymoBIOMICS™, Zymo Research) was utilized. The mock community DNA was extracted using 75 µl per preparation, following the manufacturer’s recommendations and employing the same protocol as for the other samples.

### 16S rRNA Gene Amplification and Sequencing

The V3V4 hypervariable region of the bacterial 16S rRNA gene was amplified using a two-stage polymerase chain reaction (PCR) protocol, using the 341F and 806R primers (24) and a protocol described previously (21). The final PCR libraries were pooled and sequenced using a 15% phiX spike-in on an Illumina MiSeq sequencer with V3 chemistry for 2×300 base paired-end reads. This sequencing was performed at the Genomics and Microbiome Core Facility (GMCF; Rush University, IL, USA). The raw sequences were deposited in NCBI PRJNA1107792.

### Raw Data Processing

Quality control of the raw paired-end reads was performed using FastQC v0.11.9 tool, with the rest of the analysis conducted using R 4.1.0 in RStudio 2023.6.0.421. Libraries were processes using DADA2 (25) following the recommended workflow, with forward and reverse reads trimmed to 240bp and 215bp, merging with a minimum overlap of 8 bp, and taxonomic classification against SILVA v138.1 (26). ASVs that were not classified at the phylum level or were not assigned to bacteria were excluded from further analysis.

### Data and Statistical Analysis

Data management and organization were conducted using the tidyverse v2.0.0 package (27), and results were plotted using ggplot2 v3.4.2 (28). α and β-diversity were calculated using vegan v2.6.4, mia v1.6.0 (29), and phyloseq v1.40.2 (30). We used Bray-Curtis index to create a dissimilarity matrix later used for Non-metric Multidimensional Scaling (NMDS) plots, Variation Partition Analysis (VPA), and diversity comparison. The matrix was created using the “avgdist” function from the vegan package with the dataset subsampled to 14500 across 1000 iterations for robustness. A similar method was used to calculate a mean for the α-diversity indexes using “estimateRichness” from the mia package. To statistically validate the differences observed in microbial communities, we applied Two-way PERMANOVA. Additionally, we use ARTool (31, 32) to compare differences in Bray-Curtis dissimilarity and α-diversity among samples. To identify significant taxonomic variations in relative abundance across different tissues and bloom stages, we analysed taxa with a maximum median relative abundance of 1% in the monthly analysis and 3% in the tissue-based analysis. The elevated threshold for tissues aimed to minimize the influence of bell-only taxa, as it was significantly richer than the other tissues. Taxa were further analysed only if they had a p-value below 0.05 in both the Kruskal-Wallis test and subsequent Wilcoxon test.

### Comparing *R. nomadica* microbiomes with other jellyfish and water samples

In addition to our primary dataset, we analyzed microbiome data from 9 other jellyfish species (Table S1), and from water samples collected monthly at two sites in the Eastern Mediterranean. The jellyfish microbiomes were from *Rhopilema esculentum, Nemopilema nomurai, Rhizostoma pulmo, Cassiopea xamachana, Mastigias papua, Aurelia coerulea, Cyanea nozakii, and Tripedalia cystophora*(33–36). Raw sequencing reads for these datasets were downloaded from NCBI. The water samples were collected during the SoMMoS cruises (Southeastern Mediterranean Monthly cruise Series, (20) from two sites, one at the edge of the continental shelf (surface samples only) and one from open water (2.5m-1450m depths, Table S2, (20). Two size fractions were analyzed – larger than 11 micron and 5-0.22 micron (37). The full dataset from these cruises is currently being analyzed, and we focus here only on those that are identical to the ASV sequences from jellyfish. The 16S rRNA datasets from the additional jellyfish and water samples were previously sequenced using different primer sets spanning different regions of the 16S rRNA gene (V3V4, V4V5, or V5V6). To enable a direct comparison between these datasets, all samples were re-analyzed using the same pipeline as described above, followed by a genus-level comparison (21). ASVs sequences comparisons were conducted using the Nucleotide-Nucleotide BLAST 2.12.0+ (BLASTn) algorithm (38) on a Linux platform. Only samples with overlapping V4 region were compared. Mean relative abundance for each ASVs or genus was calculated to get a representative dataset for each jellyfish species. An additional analysis was dedicated to the jellyfish species *Cotylorhiza tuberculata*, for which the raw sequence data were derived from metagenomic samples and therefore did not conform to the standard 16S rRNA gene sequencing pipeline. For this species, we used BLAST with the sequences of the top 25 ASVs from *R. nomadica* as query against the SRA database.

## RESULTS

### The *R. nomadica* microbiome differs between tissues and over time

First, we aimed to examine whether *R. nomadica* microbiome differs between jellyfish tissues and changes across different stages of the bloom (Fig. 2). In agreement with a previous, smaller, study, the bell and gonads hosted distinct communities (PERMANOVA, p<0.01), while the gastrovascular canals (GVC) and tentacles mostly overlapped (PERMANOVA, p>0.5, Fig. 2A, (21)). However, clear temporal differences were also observed (PERMDISP2 p-value < 0.001, Fig. 2B). Indeed, Variation Partitioning Analysis (VPA) analysis showed that tissue type and month of bloom are the most influential factors on the variation between samples, explaining 19% and 11% of the variation, respectively (Fig. 2C). When repeated for individual tissues separately, the VPA revealed even clearer temporal changes in microbiome structure, accounting for 17-24% of the variations (p value < 0.003, Fig. S1). Here, again, the bell and gonad communities varied significantly between months (PERMANOVA p value < 0.01), while the GVC and tentacles differed only between July (towards the end of the bloom) and the other months (PERMANOVA p value < 0.001, Fig. 2D). Other factors such as the size of the jellyfish or its sex did not significantly affect the microbial population structure (Fig. 2C).

**FIG 2.**
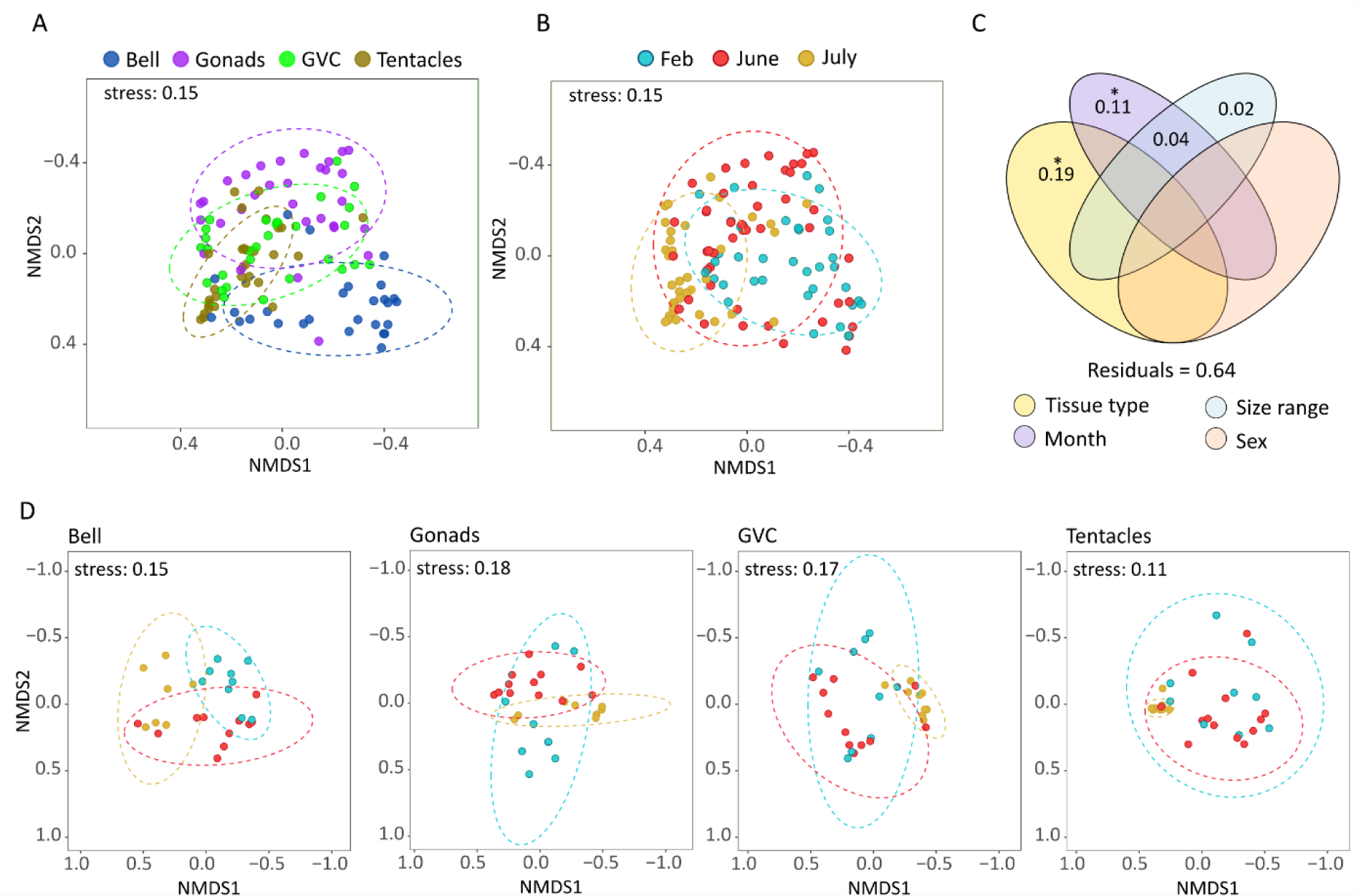
Variation in the microbiome composition. A, B) Nonmetric multidimensional scaling (NMDS) analysis for all samples, colored by tissue type (A) or month of sampling (B). Additional panels representing the third dimension can be found in Fig. S1C) VPA analysis performed on all data. Asterisks represent statistical significance p<0.001 as estimated using CCA analysis. Colors represent different factors that were tested. The total percentage of variance explained by the factors is calculated as 1 minus the residuals. D) NMDS analysis for each tissue separately. Tissue type is specified on top of each panel, the colors represent different months as specified in panel B.

Notably, the samples from July formed tighter clusters in the NMDS ordination that those from other months, particularly in the tentacles and the GVC, suggesting more uniform communities across individual jellyfish towards the end of the bloom (Fig. 2B, D, Fig. S2). Taken together, these results reveal distinct patterns within *R. nomadica* across different tissues and bloom stages, with the most pronounced shifts occurring in July.

### Key Bacteria (ASVs) Shaping Variability Across Tissues and Bloom Stages

Next, we aimed to identify the bacterial taxa responsible for the observed dissimilarities between tissues and months. A relatively small group of ASVs dominated the entire jellyfish microbial community, with 68 ASVs representing taxa with a relative abundance exceeding 1%. Notably, the bell tissue contributes to most of these ASVs making it the most diverse tissue (Fig. 3A, Fig. S3). The differences between tissues and month of sampling were primarily associated with six types of bacteria, which, while present to some extent in all tissues, show clear temporal patterns (Fig. 3B). The most relatively abundant ASV, an *Endozoicomonas* (ASV1), was associated primarily with the tentacles, and to a lesser extent with the bell and GVC. ASV1 was also highly seasonal, peaking in all tissues during July, when it comprised, on average, more than 89% of the tentacle community. The second most abundant ASV was an unclassified Rickettsiales (ASV2), with high relative abundance mostly in the gonads and GVC, also peaking during July. Other noteworthy associations were between an ASV belonging to the Entomoplasmatales (ASV3) with the tentacles and GVC, mostly during February and June; a member of the Simkaniaceae (ASV4) with the gonads during June, and a member of the *Bacteroides* (ASV6) with the bell primarily in February. The gonads were the only tissue that had a high relative abundance of *Mycoplasma (*ASV11/21) (Fig. 3A-C). Finally, a *Tenacibaculum* (ASV5) was clearly associated with the winter bloom, without any clear tissue pattern.

**FIG 3.**
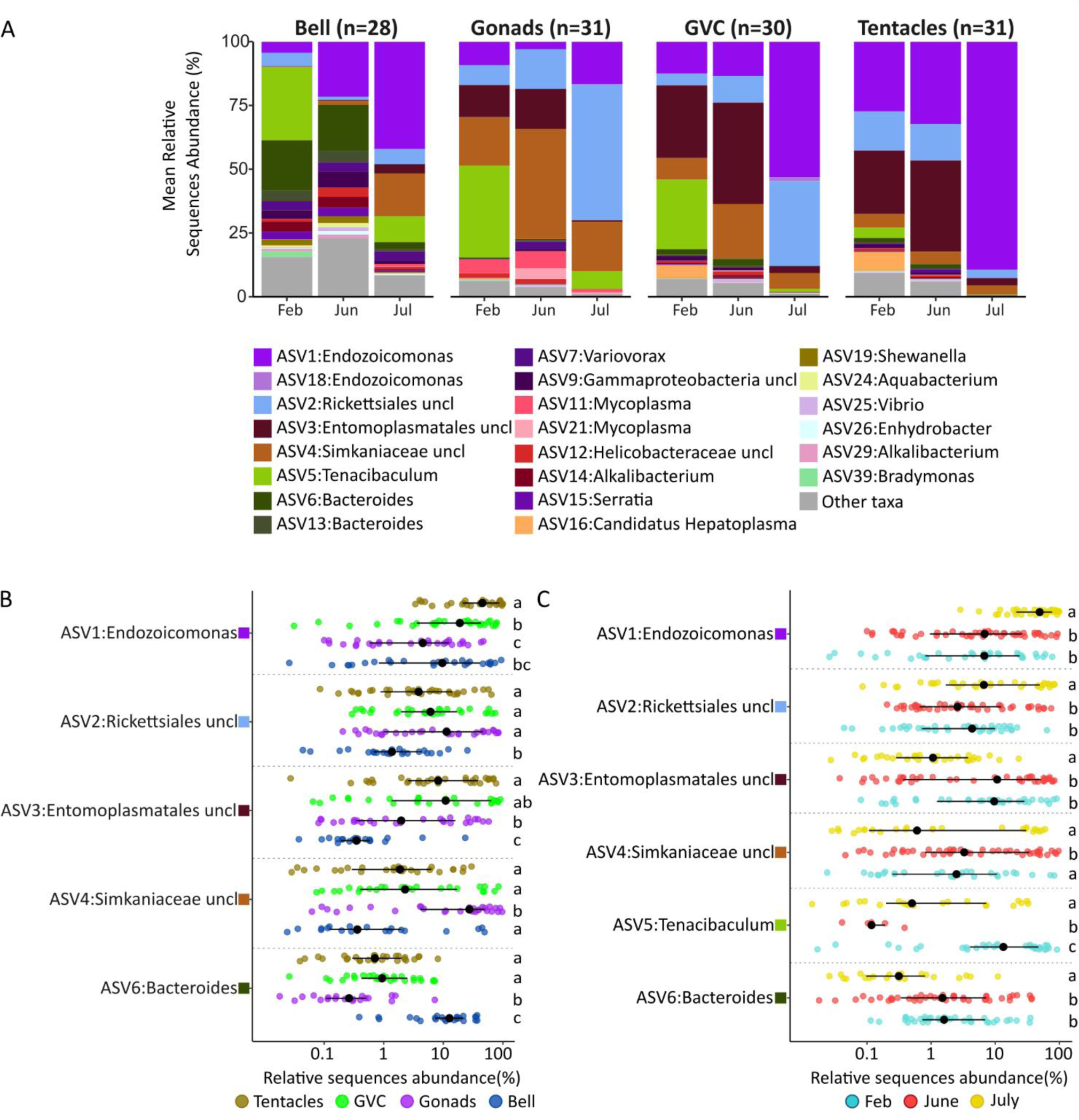
Jellyfish bacteria composition analysis. A) Mean relative abundance analysis for tissues across bloom months. Type of tissue is specified above each graph. ASVs belonging to the same genus are color coded with similar colors and grouped together, otherwise taxa are ordered by mean relative abundance. Each ASV is classified to the highest level of phylogenetic identification achieved. The data for all individual samples are shown in Fig. S4. C) Taxa with significant differences between tissues (B) or months (C). Colors are the same as those used in Fig. 2. Letters represent significant groups found in with p < 0.05 as determined by pairwise comparisons-Wilcoxon tests following a Kruskal-Wallis test.

### Microbial Alterations during Jellyfish Bloom Decline

To investigate the link between microbial alterations and the decline of the bloom, we concentrated on two potential scenarios: the emergence of pathogenic bacteria or a significant shift in microbial community structure leading to an imbalanced microbiome. Both scenarios could potentially disrupt the physiology of the jellyfish, thereby affecting the bloom.

First, we sought to examine how the health of jellyfish influences the diversity and composition of their microbiome. We focused on seven individuals exhibiting clear signs of damage, including major lesions, holes, unusual morphology, or a heavily cracked bell. We obtained samples from these damaged areas, as well as from visibly normal regions from the same individuals. Our analysis revealed no notable distinction between the microbiota of healthy and unhealthy jellyfish. The monthly clustering patterns remained consistent, with samples from both healthy and unhealthy individuals interspersed (Fig. S5). We found no specific taxa that were exclusive to either group. This observation remained consistent even when we focused our analysis on comparing damaged and undamaged areas within the same individual jellyfish, where we still found no discernible patterns (Fig. S5)

Next, we examined the α-diversity across various stages of the bloom considering that shifts in microbial diversity could critically impact the functionality and resilience of the host. The Shannon diversity during July (late-stage bloom) was notably lower than in June (early-stage bloom) or February (winter bloom) (Aligned Rank Transform (ART) ANOVA test, p-value<0.001, Fig. 4A). We also examined the Bray-Curtis distances to explore differences between bacterial communities of each month. Our results indicated that the communities in July were significantly more homogeneous compared to those from other months (ART ANOVA test, p-value<0.001, Fig. 4B). Upon further examination, we found that in July, the bell and tentacles exhibited lower diversity, while more uniform communities were observed in the gonads, tentacles, and GVC (Fig. S2).

**FIG 4.**
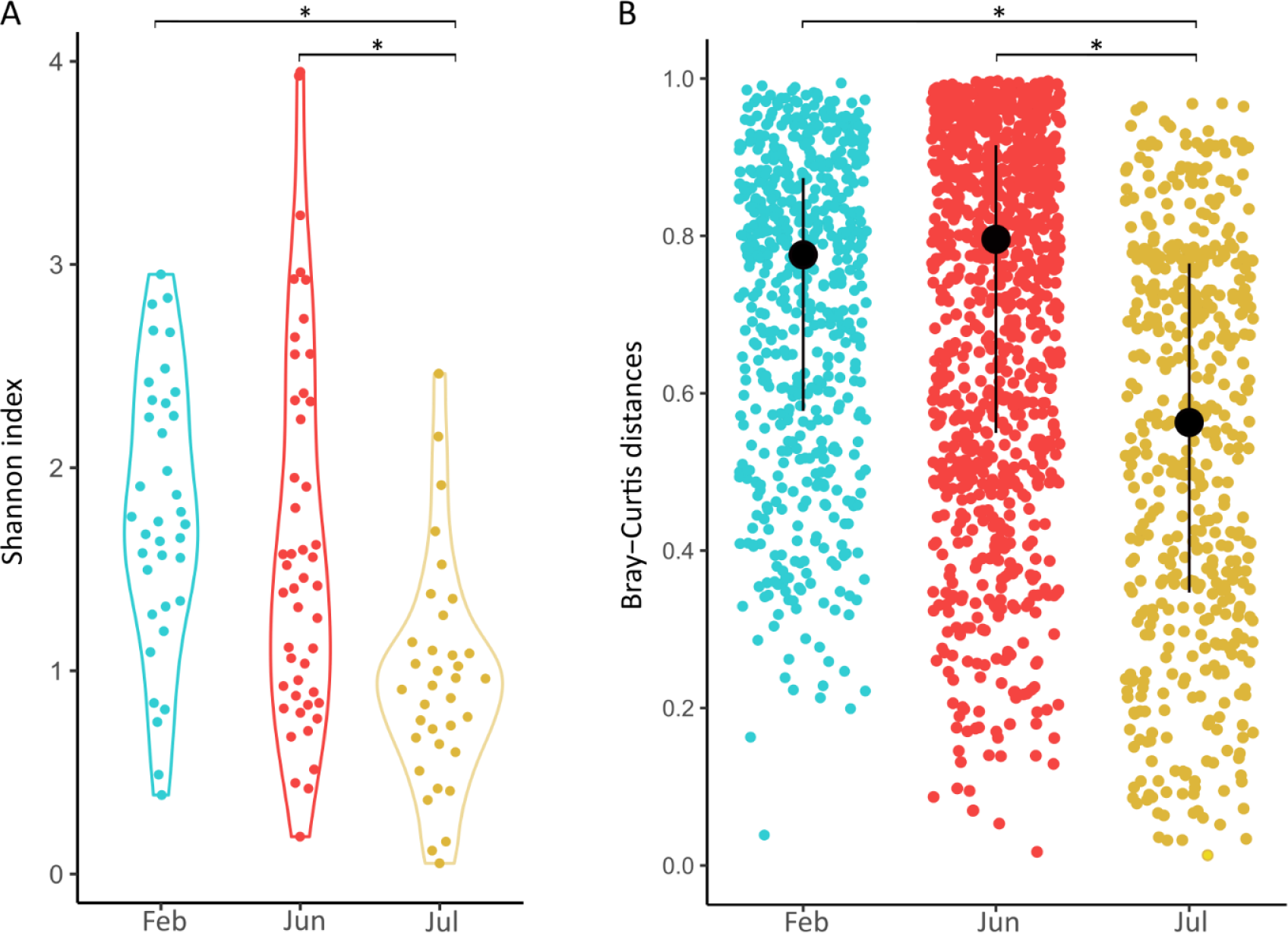
Variation in bacterial community diversity over different bloom events. A) Shannon diversity of individual samples. B) Bray Curtis dissimilarity between samples from the same month. Asterisks represent statistically significant differences (Aligned Rank Transform (ART) ANOVA test, p-value<0.001)

Overall, no specific taxa or patterns could differentiate healthy and unhealthy jellyfish, yet a significant decline in microbiome diversity was observed during the late bloom stage.

### Microbial Connectivity of *R. nomadica* with its Environment and Other Jellyfish Species

In this section, we focus on two questions. First, we ask whether a ‘core’ microbiome exists across jellyfish species. This could indicate whether certain clades or strains are essential across species. Second, we ask whether ASVs found in *R. nomadica* are also found in the water surrounding the jellyfish, or in a “baseline” time-series where jellyfish were not observed during sampling. With these data we can begin to ask how the jellyfish acquire their microbiome – whether they are colonized by bacteria commonly found in the surrounding water.

When the most dominant bacterial community members across all jellyfish were examined, no single ASV (Fig. 5A) or even genus (Fig. 5B) was found, thus no clear ‘core’ could be identified. Additionally, there was no clear pattern that could be attributed to the phylogeny of the jellyfish species, i.e. there were no ASVs or genera that are found consistently in a subset of closely related jellyfish. Some genera, such as *Endozoicomonas*, *Spiroplasma*, *Mycoplasma* and *Rasltonia*, appear in over half of the jellyfish (5-6/9), where they often exhibit high relative abundance (Fig. 5B). In contrast, while *Vibrio* was the most prevalent genus, appearing in 8/9 jellyfish, it is found in very low relative abundance in all except *Aurelia coerulea* (Fig. 5B).

**FIG 5.**
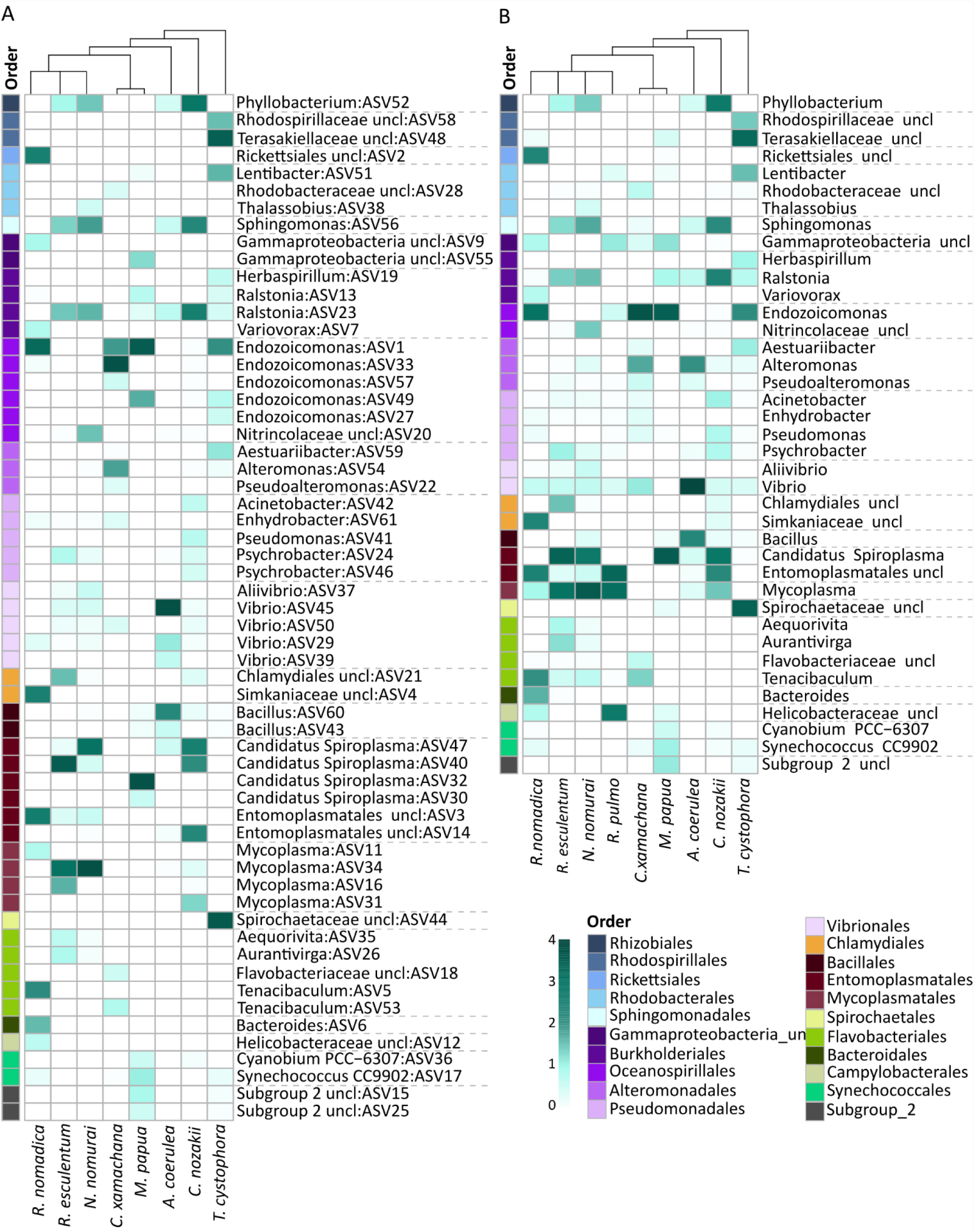
Comparative analysis of microbial populations in different jellyfish species. A) ASV-level relative abundance. The heatmap displays the top 10 ASVs for each jellyfish species after a 100% sequence similarity BLAST match, yielding 59 unique ASVs defined based on part of the conserved V4 region. When more than one ASV has identical V4 regions (but, for example, differ in other sequenced regions) they are combined to include a single representative ASV from each species. B) Genus-level relative abundance heatmap. Displays the aggregation of all ASVs at the genus level identified in panel A and incorporating additional *Rhopilema pulmo* sequences from the V5V6 16S region (which do not overlap with the V4 region and thus could not be included in the ASV comparison in panel A). Relative abundances are represented through a color gradient, normalized using log(1+x) to accommodate both low and high values. A phylogenetic tree above the heatmaps orders jellyfish species by their evolutionary relationships, aiding in the detection of microbial composition patterns.

Within the predominant genera, specific ASVs of *Endozoicomonas* (ASV1) and Ralstonia (ASV23) were identified across diverse geographical locations, highlighting a potential widespread distribution. *R. nomadica* from the Eastern Mediterranean shared an identical *Endozoicomonas* ASV with *Cassiopea xamachana* from Nichupté Lagoon in Mexico (maintained in the lab), and with *Mastigias papua* and *Tripedalia cystophora* from Indonesian marine lakes (Fig. S6). This indicates a shared microbial community among geographically distant species. In contrast, ASVs from other genera showed associations with specific geographical locations. For example, the ASVs identified in the four jellyfish species (*Rhopilema esculentum*, *Nemopilema nomurai*, *Cyanea nozakii*, and *Aurelia coerulea*) from the Yellow Sea appeared to correlate with their respective habitats, suggesting a degree of localization (Fig. S6).

The microbial diversity in the seawater collected from within the jellyfish bloom was higher than that of the jellyfish tissues and was clearly different from it in an NMDS analysis (Fig.S3). From the 25 most abundant ASVs in the *R. nomadica* microbiome, 8 were shared with the water surrounding the swarm, and 5 were shared with the water collected from the “baseline” locations (Fig. 6). Most of these shared ASVs were rare (<0.1% average relative abundance) in both jellyfish tissues and seawater, with the exception of *Synechococcus* CC9902 (ASV8), which is widely present in seawater and formed ∼20% of the reads within the sampled jellyfish swarms. It is noteworthy that the three ASVs with the highest mean relative abundance in the jellyfish tissues were identified in the water within the swarm but not in the baseline sites (*Endozoicomonas* ASV1, Rickettsiales (ASV2) and unclassified Entomoplasmatales (ASV3). The unclassified Entomoplasmatales (ASV3), which closely matches (99.5%) a complete 16S sequence from *C. tuberculata* (39), has been further identified as *Spiroplasma* and will henceforth be referred to as *Spiroplasma*.

**FIG 6.**
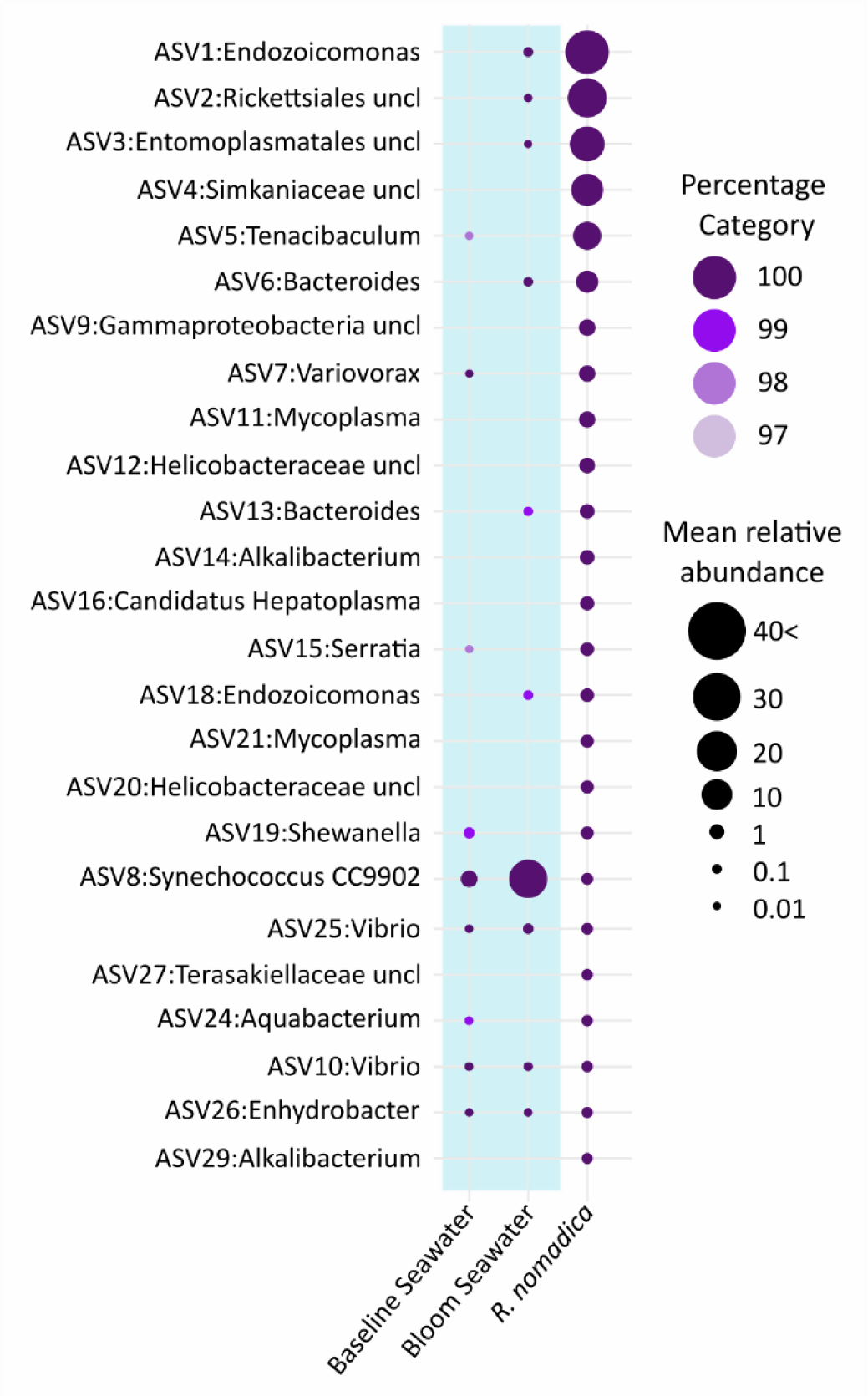
Comparative analysis of top *R. nomadica* ASVs against seawater from different locations. The comparison was done on ASV level. The size of each circle indicates the mean relative abundance of the corresponding ASV, while the color denotes the percentage identity among the ASVs examined. The blue background represents the two seawater types: water from the bloom and baseline water collected from a remote location.

## DISCUSSION

In this study, we examine the microbiome dynamics of *R. nomadica* across various tissues and bloom stages. We observed a decrease in microbial diversity as the bloom progressed, with taxa like *Endozoicomonas* and Rickettsiales becoming dominant in specific tissues. Extending our analysis to other jellyfish species revealed that although certain clades, and even specific ASVs, such as *Endozoicomonas*, are common, they are not universal, indicating a lack of a “core jellyfish microbiome.” Interestingly, certain ASVs, including the three most abundant, were present only in the bloom water and were absent in distance water where jellyfish were not present. The discussion below will further explore these changes and their implications for jellyfish health and ecosystem dynamics.

### Temporal Microbial Shifts in *R. nomadica* Are Characterized by Reduced Communities Diversity

Throughout the months of February, June, and July, we observed significant shifts in the microbial communities of *R. nomadic*a, especially noticeable between the early bloom in June and the late bloom in July. The late bloom stages featured less diverse, more homogenous bacterial communities across all tissues. These shifts were predominantly driven by two taxa*, Endozoicomonas* (ASV1) and Unclassified Rickettsiales (ASV2), which were present in over 90% of our samples and suggest a pivotal role in host-microbe interactions (40–43). However, this dominance may lead to an imbalanced microbial community with potential negative impacts on the host (40, 44, 45). Additionally, these changes could correlate with physiological changes in the jellyfish, such as the degradation of feeding structures, affecting food intake and thus the microbiome (46). These observations indicate that changes in microbial dominance and diversity might not necessarily reflect the health status of jellyfish but may be part of normal developmental processes the jellyfish go through as the blooms evolve. This raises questions about the dynamic roles these bacteria play in jellyfish, particularly how their interactions with the host might change during different bloom stages.

*Endozoicomonas* are known for their interactions with multiple marine hosts, particularly cnidarians, and have been extensively studied in corals. They often appearing in high abundance, forming aggregates, and important to coral health and resilience (47–52). However, their relationship may not be solely mutualistic, as indicated by genomic traits like eukaryote-like proteins, secretion systems, and antioxidant properties, which suggest a can also be associated with pathogenic lifestyle (53, 54). This implies a complex relationship where *Endozoicomonas* could occupy various niches across the symbiotic spectrum, including potential negative effects on health and fitness. Different *Endozoicomonas* strains can have varied roles, with specific clades linked to certain cnidarian families, and others found across diverse species (53, 55). Their responses to environmental stressors, such as heat, also vary, with some studies linking a decrease in *Endozoicomonas* abundance to coral bleaching events, while others report an increase or unchanged composition (50, 56–58).

The role of Unclassified *Rickettsiales* (ASV2) in jellyfish remains speculative, given their classification only to the order level in *R. nomadica*. Generally, *Rickettsiales* are considered obligate intracellular bacteria, often host-specific and potentially pathogenic (59, 60), with occurrences reported across various marine hosts, including bivalves and corals (61, 62). For example, in the coral *Acropora cervicornis*, a low abundance of *Rickettsiales* spp (*Candidatus Aquarickettsia*) was tolerant by the coral, while high abundance was hypothesized to compromise host health by depleting it nutrients and energy Although deeper taxonomy identification is needed for the *Rickettsiales* spp. here, it is possible that the shift in *Rickettsiales* during July bloom may trigger disease processes and therefore possibly indicate unhealthy/ageing jellyfish. Interestingly, the *Rickettsiales* sequence in *R. nomadica* did not match any other jellyfish, suggesting a species-specific relationship.

The observed increase in *Endozoicomonas* and *Rickettsiales* during the late bloom stage may suggest a transition from a neutral or beneficial symbiosis to a potentially harmful one. This microbial shift, possibly influenced by environmental conditions or physiological changes within the jellyfish, coincides with the decline of the bloom, suggesting an adverse interaction. However, fully understanding these dynamics requires further research.

### *Tenacibaculum* spp. Dominate *R. nomadica* Winter Blooms

Unlike the well-documented summer blooms, the dynamics, and connections of winter blooms to their summer counterparts remain less understood. Our analysis over consecutive winters (2020 and 2021) revealed a consistent winter bacterial community, which is different from the summer one. Genetic analysis show that the winter and summer blooms appear to be part of the same genetically diverse population indicated by high gene flow and lack of genetic differentiation between seasons and months This suggests that the microbial differences between seasons are driven by environmental conditions or jellyfish physiology rather than distinct cryptic species (64, 65).

The winter community has a high relative abundance of *Tenacibaculum* spp (ASV5), which is mostly absent in early summer and sporadically observed in late summer. *Tenacibaculum* species, part of the Flavobacteriaceae family, show a range of interactions with marine hosts, from symbiosis to pathogenicity. Although their exact role in jellyfish is not fully understood, they are observed across various jellyfish species (Fig. 5B (13)), with different ASVs present in each (Fig. 5A). *In Cotylorhiza tuberculata*, *Tenacibaculum*-like bacteria within the gastric filaments may enhance carbon and energy transfer, suggesting a symbiotic relationship. Their genome encodes pathways for glycolysis, the Krebs cycle, and synthesis of essential amino acids and vitamins such as thiamine (B1), biotin (B7), and riboflavin (B2), providing significant metabolic benefits to the host (38, 65). Additionally, species like *Pelagia noctiluca* and *Phialella quadrata* are potential reservoirs for *Tenacibaculum maritimum*, a known pathogen of farmed fish (66, 67). However, in *R. nomadica*, the presence of *Tenacibaculum* appears non-obligatory and possibly influenced by environmental factors.

### Key Microbial Taxa in *R. nomadica* – Putative Roles and Cross-Species Presence

Our analysis reveals that the *R. nomadica* microbiome is dominated by a few bacterial taxa with high relative abundance and prevalence, consistent with other jellyfish species (33, 36, 39, 68). We investigated whether key taxa are shared across jellyfish with different phylogenetic and geographic backgrounds but found no consistent ‘core’ microbiota at the ASV or genus level (Fig. 5). Additionally, phylogenetic classification did not show any distinct patterns indicative of of phylosymbiosis, aligning with studies suggesting that shared bacteria among cnidarians and marine invertebrates often reflect environmental rather than phylogenetic similarities (52, 55, 69). In this regard, we observed that certain ASVs were exclusive to four jellyfish species from the Northern Yellow Sea (33). However, these four jellyfish were from a single study, and thus further work is needed to determine if the sampling location influences the microbiome composition of different jellyfish species.

Despite the absence of a clear shared core microbiome our analysis identified several bacterial taxa common to various jellyfish species and locations. Notably, an *Endozoicomonas* ASV (ASV1) and a *Ralstonia* ASV (ASV23) were widespread across diverse environments ASV1 was particularly abundant in *R. nomadica* from the Eastern Mediterranean and was also highly relative abundant in other species like *M. papua* and *T. cystophora* in Indonesian Marine lakes, and *C. xamachana* from Mexico’s Nichupté Lagoon, which was cultured in a lab. According to McCauley et al. (2023), *Endozoicomonas* ASVs are divided into generalist and host-specific groups, with some clades like those in Atlantic Rhizostomeae being highly host-specific. Our *Endozoicomonas* ASV1 matches ASV59, 97, 1987, 2517, and 96488, which McCauley et al. classified as specific to this group. This supports the association of the clade with jellyfish but also suggests its adaptability beyond previously observed geographic constraints.

Focusing only on ASV-level bacteria might miss broader patterns evident at higher taxonomic levels, leading us to also examine genera common among jellyfish. Notably, *Mycoplasma*, *Spiroplasma*, and *Vibrio* emerged as highly prevalent taxa. This prevalence warrants further exploration into their roles in jellyfish, which might explain their widespread presence across different geographical and phylogenetic backgrounds.

While the specific functions of *Spiroplasma* in jellyfish remain unclear, they are recognized for their intracellular lifestyle and defensive roles in host interactions (70, 71). Observations in *C. tuberculata* jellyfish suggest that they have a small genome and predicted anaerobic metabolism, hinting at a commensal relationship (39). *Mycoplasma*, typically viewed as intracellular parasites, have recently been identified as part of the core microbiome in healthy cnidarian tissues (55, 72, 73), and their role may shift from beneficial to harmful due to changes in the community composition or environmental factors (6, 11). Lastly, Vibrio present an interesting case as while very prevalent among the jellyfish species, they appear in low abundance and consistently found in all our seawater samples. This finding underscoring their generalist nature and consistent with the prevalence of the Vibrionaceae Family in Anthozoa, Cubozoa, and Scyphozoa (55). We will discuss these findings further in the next section.

### How did the jellyfish get its bacteria?

Previous studies indicates that cnidarian and other invertebrate microbiomes are distinct from the surrounding seawater (13, 55, 69). However, ASVs within cnidarian microbiomes often show region-specific patterns (14, 55), hinting at potential organism-to-organism transfer. How, then, are cnidarian-associated bacteria acquired? Are they present in seawater at low abundances, possibly associated with marine particles, thus potentially “inoculating” cnidarians? Or, are they exclusively around cnidarians, transferred vertically or through contact with other cnidarians or their mucus? Our comprehensive sampling of jellyfish and surrounding waters, including a control site without jellyfish, allowed us to explore these modes of transmission. Focusing on the main *R. nomadica* ASVs, we categorized the main *R. nomadica* ASVs into three groups-1. Shared between jellyfish, bloom seawater, and ‘control’ seawater 2. Shared between jellyfish and bloom seawater 3. Found only in the jellyfish.

The first group includes marine bacteria like two *Vibrio* ASVs (ASV10 and ASV25), commonly found in the jellyfish microbiome and are likely opportunistic, adapting to various niches such as marine particles and different jellyfish species, including those in this study. *Vibrio* are known to inhabit diverse marine niches, including various cnidarian hosts (13, 55, 74, 75). Notably, *Vibrio* increase in abundance during the decay of *Aurelia aurita* jellyfish blooms, underscoring their role as opportunistic decomposers (8, 76). Additionally, *Vibrio* produce transporters that aid in nutrient uptake from decomposing jellyfish, leveraging public goods released by others and creating niche partitioning within the bacterial community (76).

The second group includes bacteria found on jellyfish and in bloom water, but absent in jellyfish-free water, featuring four primary ASVs from *R. nomadica*—*Endozoicomonas* (ASV1), unclassified Rickettsiales (ASV2), *Spiroplasma* (ASV3), and *Bacteroides* (ASV6). *Endozoicomonas* may have a free-living stage as suggested by their large genome, metabolic diversity, and ability to utilize diverse carbon sources (48, 49, 56). However, our findings, reinforced by coral studies, indicate that their presence in seawater primarily depends on proximity to hosts. For example, *Endozoicomonas* abundance decrease with increasing distance from coral reefs (77). Additionally, two generations of adult *Pocillopora acuta* displayed consistent *Endozoicomonas* ASVs, which were absent in their larvae (47). This suggests horizontal acquisition at the adult stage, emphasizing a primarily host-associated lifestyle rather than a truly free-living existence.

In contrast to *Endozoicomonas*, neither Rickettsiales—primarily intracellular pathogens or symbionts— nor *Spiroplasma* and Bacteroides, both considered anaerobic, are expected to have a free-living stage (39, 78, 79). *Spiroplasma*, identified as anaerobic fermenters in *C. tuberculata* (showing 99.53% identity with ASV3 from *R. nomadica* microbiome), were found primarily in the tentacles and gastric vascular cavity (GVC), and also in the mucus of *R. pulmo*(39, 80). Similarly, Bacteroides, typically studied in mammalian gastrointestinal tracts as obligate anaerobes, has limited known environmental survival abilities (81). Here, they were mostly found in the bell of *R. nomadica*, and we speculate that they may be associated with the bell mucus layer, akin to their role as mucosal bacteria in animal guts (82). However, the identification of putative anaerobic organisms in the jellyfish microbiome raises the question of whether and where these animals have anaerobic environments. (20). Their presence in the water surrounding the jellyfish bloom could be explained by the shedding of tissue or mucus, as suggested in corals (77).

The final group of ASVs, found exclusively on the jellyfish, includes unclassified Simkaniaceae (ASV4) and *Mycoplasma* (ASV11). These were primarily identified in the gonads of *R. nomadica*, raising the possibility for vertical transmission. Similarly, in *Pocillopora acuta*, a member of the *Simkania* (a *Simkaniaceae* genus) appeared not only in adults but consistently across larval samples, indicating vertical transmission in asexually reproducing larvae (47). Likewise, in *Aurelia aurita*, the presence of *Mycoplasma* in adults and various developmental stages also supports the likelihood of vertical transmission among these cnidarian species (75, 83).

In addition to inoculation by waterborne bacteria and vertical transmission, food-borne bacteria (e.g. associated with copepods or other zooplankton, (46) could also be a source for jellyfish-associated communities. Intriguingly, *Simkania*-like cells, corresponding to our ASV4 unclassified Simkaniaceae, were found inside eukaryotic ciliates in the gastric cavity of *C. tuberculate* (39), suggesting potential tri-partite symbioses (84). Investigating environmental conditions within jellyfish niches, like oxygen levels in mucus, and conducting broader sampling of surrounding waters including prey- and other eukaryote-associated bacteria, could clarify these transmission hypotheses. Additionally, studying other jellyfish life cycle stages, such as polyps and ephyrae 82, 84, 85), might provide deeper insights into the mechanisms governing jellyfish-associated communities and how these microbiomes influence jellyfish fitness.

## ACKNOWLEDGMENTS

We thank the crew of the Mediterranean Explorer for assistance during the jellyfish sampling, all of the students from the Leon H. Charney School of Marine Sciences, University of Haifa, who helped to collect the jellyfish, and Jennifer Hennenfeind for the DNA extractions of the SoMMoS cruise samples.

## DATA AVAILABILITY STATEMENT

Raw 16S sequencing data from this study are available in the NCBI Sequence Read Archive under Accession Number PRJNA1107792. The analysis code is accessible on GitHub at https://github.com/NogaBarak/R.nomadica_microbiome. Samples metadata and ASV table can be found in Tables S3 and S4.

